# Automated Joint Space Detection Improves Bone Segmentation Accuracy

**DOI:** 10.1101/2025.04.30.651481

**Authors:** H. Mark Kenney, Daniel Lichau, Rémi Blanc, Lindsay Schnur, Richard D. Bell, Christopher T. Ritchlin, Edward M. Schwarz, Hani A. Awad, Ronald W. Wood

**Affiliations:** Center for Musculoskeletal Research, University of Rochester Medical Center, 601 Elmwood Ave, Rochester, NY, USA, 14642; Department of Pathology & Laboratory Medicine, University of Rochester Medical Center, 601 Elmwood Ave, Rochester, NY, USA, 14642; Department of Medicine, University of Rochester Medical Center, 601 Elmwood Ave, Rochester, NY, USA, 14642; Thermo Fisher Scientific, 3 Impasse Rudolf Diesel, Merignac, France, 33700; Arthritis and Tissue Degeneration Program, Department of Research, Hospital for Special Surgery, 535 E71st St, New York, NY, USA, 10021; Department of Obstetrics and Gynecology, University of Rochester Medical Center, 601 Elmwood Ave, Rochester, NY, USA, 14642; Department of Neuroscience, University of Rochester Medical Center, 601 Elmwood Ave, Rochester, NY, USA, 14642; Department of Urology, University of Rochester Medical Center, 601 Elmwood Ave, Rochester, NY, USA, 14642; Department of Biomedical Engineering, University of Rochester Medical Center, 601 Elmwood Ave, Rochester, NY, USA, 14642; Department of Orthopaedics, University of Rochester Medical Center, 601 Elmwood Ave, Rochester, NY, USA, 14642

**Keywords:** Micro-CT, segmentation, image analysis, image processing, deep learning, arthritis

## Abstract

Quantitative description of complex anatomical structures remains challenging due to the expertise necessary for manual segmentation, labor, and interobserver variability. To overcome this, automated detection of specific landmarks can be provided by digital image analysis techniques including deep learning (DL) models. To this end, we undertook supervised automated analysis of micro-computed tomography (micro-CT) datasets of murine hindpaws and forepaws (30-33 bones). Advancing beyond previously published semi-automated (SA) marker-based watershed algorithms, we added structure enhancement, tensor voting, and output dilation to identify joint spaces. Segmentation was enhanced by the use of a DL joint space prediction model (3D U-Net architecture, ResNet-18 backbone) using wild-type (WT) hindpaw labels as ground truth. Prediction was then extended to hindpaws (52.4% in test group) and forepaws from WT and tumor necrosis factor transgenic (TNF-Tg) mice with inflammatory-erosive arthritis of both sexes across age. Segmentation accuracy improved dramatically using the DL methodology (WT male: SA 79.39±5.73% vs DL 98.16±1.47%, *p<0.0001*; WT female: SA 79.16±4.84% vs DL 99.19±1.63%, *p<0.0001*). Accuracy declined with increased disease severity and age in TNF-Tg mice (TNF-Tg male 93.54±4.73%, female 91.81±2.80%, *p*≤*0.01*). Subsequent testing in forepaws also displayed progressive reduction in accuracy with increasing arthritic severity (i.e., WT 87.29±2.07%, TNF-Tg male 72.65±11.70%, *p<0.0001*). Overall, this supervised automated model outperforms recent SA approaches in healthy joints to enhance investigation of complex bone anatomy. Although flexible application to novel and disease-modified datasets demonstrates deprecated performance, utilization may nonetheless catalyze structure-specific segmentation model development.

## 1. Introduction

High quality image analysis not only enhances research endeavors, but also has the potential to aid clinical radiologists as they seek to detect and quantify pathologic changes, a task of paramount importance to patient care. Image analysis is a detailed sequence of procedures including feature extraction, translating an otherwise generic image into meaningful labels and then deriving quantitative metrics. Much of this process is driven by prior knowledge, such as thresholding of structures based on density or color, and then applying downstream image processing algorithms (i.e., dilation, erosion, smoothing, separation) to achieve the desired segmentation. Once optimized, the segmented images can provide inputs for supervised machine learning, including deep learning (DL) that encodes and decodes complex features using neural networks (Konnaris et al., 2022; Najjar, 2023) resulting in improved image segmentation accuracy and throughput.

Arthritis research imaging focuses on articulating surfaces between two or more bones, where methods for bone separation are critical for successful evaluation of differential pathologic processes in complex joints. Advances in image processing algorithms have demonstrated remarkable utility in increasing analytical throughput of closely adjacent carpal or tarsal bones (Kenney et al., 2022b; Sebastian et al., 2003). However, adoption has been limited by both inaccuracies requiring user intervention and difficulties in translation to distinct structures, highlighting the need for implementation of optimized workflows. These multi-step processes can benefit tremendously from discrete tools for image enhancement (i.e., bone edge detection (Besler et al., 2021)) and incorporation of automated segmentation methods for structures that theoretically emulate the shape and dimensions of even unrelated objects (i.e., vasculature (Xia et al., 2022) may resemble the thin articulations between bones). Beyond strictly morphologic operations, other studies have implemented registration-based techniques leveraging the typical reliability and consistency in anatomy for structure identification (Baiker et al., 2010). The alternative of manual inputs to generate ground truth labels is expensive and tedious but can be similarly successful, where utilization may be essential in complicated closely interlocking bones with less discrete boundaries (i.e., skull (Wang et al., 2021)). Similarly, alternative imaging approaches with multi-color/hue variability even within a discrete structure, such as magnetic resonance imaging (MRI) (Ambellan et al., 2019; Kushwaha et al., 2023; Ramos et al., 2024) or tissue histology (Bell et al., 2024), also exhibit complexity that may benefit from initial manual segmentation to guide automated processes. Together, these methods can offer additional benefits by fueling further automation, where outcomes serve as training datasets to implement DL approaches. The benefits of segmentation automation are numerous, but particularly these methods allow for detailed spatially-relevant quantitative metrics, including regional/bone-specific erosive volume changes (Brown et al., 2017; Cambre et al., 2018; Kenney et al., 2024b; Mahdi et al., 2023) as well as identification of areas with high-susceptibility to damage (Saillard et al., 2024).

Herein we build upon established semi-automated (SA) murine hindpaw segmentation methods (Kenney et al., 2022b) with improvements in image processing algorithms in combination with ground truth bone segmentations (Kenney et al., 2024a) to train DL models for joint space detection. We also demonstrate the potential for implementation of these technical advanced in novel structures, including forepaws and hindpaws with severe erosive arthritis. The associated methods developed in Amira software (Supplementary Files) and the relevant datasets are provided publicly to support adoption and collaboration in further research endeavors (Kenney et al., 2024a; Kenney et al., 2025).

## 2. Methods

### 2.1 Animal Models

All animal experiments were performed on IACUC protocols approved by the University Committee for Animal Resources at the University of Rochester Medical Center. The mice were housed in an AAALAC accredited vivarium. A total of 19 mice were used for the described experiments, including n=4 wild-type (WT) male, n=4 WT female, n=4 tumor necrosis factor transgenic (TNF-Tg) male, and n=7 TNF-Tg female with longitudinal monthly assessment. TNF-Tg mice (3647 line, C57BL/6 genetic background (Keffer et al., 1991)) were initially obtained from Dr. George Kollias with continued maintenance at the University of Rochester. TNF-Tg mice develop chronic, progressive, and spontaneous inflammatory-erosive arthritis (Li and Schwarz, 2003) with more rapid onset of articular and extra-articular manifestations in female mice leading to early mortality at approximately 5-6-months (Bell et al., 2019). Thus, additional mice were allocated to the TNF-Tg female cohort. Indicated in previous descriptions of this study cohort (Kenney et al., 2024b), n=2 TNF-Tg females died prior to completion of the study, n=1 before 4-months and n=1 before 5-months. As the TNF-Tg mice exhibit well-established asymmetric disease progression in hindpaws (Kenney et al., 2022a; Li et al., 2011), individual limbs were considered the unit of measure (2 forepaws and 2 hindpaws per animal).

### 2.2 Micro-CT Image Collection

Micro-CT datasets were collected as previously described (Kenney et al., 2022b; Kenney et al., 2024b). Briefly, the mice were placed in a Derlin plastic and clear acrylic tube with 1-3% isoflurane anesthesia for imaging with a VivaCT 40 (Scanco Medical, Bassersdorf, Switzerland) micro-CT using the following parameters: 55 kV, 145 μA, 300 ms integration time, 2048 x 2048 pixels, 1000 projections over 180°, resolution 17.5 μm isotropic voxels. The hindpaws and forepaws were taped together for stabilization during imaging sessions. Each dataset was collected in approximately 30-45 min (60-90 min total) with hindpaw and forepaw data derived from the same animals and timepoints. The mice were evaluated at monthly intervals starting at 2-months of age until 5-months (females, TNF-Tg with early mortality (Bell et al., 2019)) or 8-months (males). All or portions of the hindpaw data utilized for this study was previously published (WT: (Kenney et al., 2022b; Kenney et al., 2023); WT and TNF-Tg:(Kenney et al., 2024b)) and are publicly available (Kenney et al., 2024a). The forepaw data was made publicly available for the purposes of this study (Kenney et al., 2025).

### 2.3 Joint Space Segmentation Algorithm with Deep Learning Facilitation

We recently developed a high-throughput image processing algorithm to segment the individual bones of the complex hindpaw in mice (30-31 bones) (Kenney et al., 2022b), which provided a framework to investigate individual biomarkers of inflammatory-erosive arthritis progression in a practical manner using Amira software (ThermoFisher Scientific, FEI, Hillsboro, OR, USA) (Kenney et al., 2024b). This baseline SA segmentation method utilizes a marker-based watershed algorithm (Beucher and Meyer, 1992) for separation at bone boundaries. The marker-based watershed algorithm separates different objects in an image by treating pixel values as local topography based on user-defined markers. These bone-specific markers were generated in a semi-automated fashion through a variety of image processing steps, including black top-hat (BTH) in an effort to highlight local regions of large density changes, such as bone edges and articulations. Together, this approach created an eroded version of each individual bone that were then expanded to bone borders through application of a binary boundary mask.

While the watershed method improved upon previous utilization of manual contouring, the accuracy (bones segmented correctly / total bones) was approximately 80% per dataset due to low contrast noise leading to bridged joint regions (over-connecting bones; 2+ bones as 1 material) or edge misidentification (over-splitting bones; 1 bone as 2+ materials). Thus, the SA marker generation for the watershed approach required consistent and frequent manual correction procedures (Kenney et al., 2022b) to eventually develop a resource of gold-standard labels (Kenney et al., 2024a; Kenney et al., 2024b) to guide further algorithmic advancements.

In this study, we build upon the foundation of our prior work by combining our BTH approach with a combination of structure enhancement (Frangi et al., 1998) to capture joint areas that emulate dark thin surfaces (akin to vessels) and tensor voting (Martinez-Sanchez et al., 2014) to reinforce joint space continuity by limiting membrane gaps. Together, these approaches fortify the joint spaces for bone separation to limit the segmentation leakage between adjacent bones that then generate over-connection errors as the watershed algorithm propagates across multiple bones. In addition to these image processing enhancements, we incorporated a DL component utilizing the gold-standard bone segmentations as ground-truth labels (Kenney et al., 2024a; Kenney et al., 2024b) for training of joint space identity as the peri-articular negative space, as described below. Thus, the final method (Supplementary Files) developed in Amira (v2022.2) utilized modules incorporating BTH (Kenney et al., 2022b), Structure Enhancement Filter (SEF) (Frangi et al., 1998), Membrane Enhancement Filter (MEF; based on tensor voting) (Martinez-Sanchez et al., 2014), DL of joint space, and further dilation with a half-kernel size of 2 to separate original micro-CT datasets into bone-specific labels.

The development of the DL model for bone joint prediction was based on the architecture of a 3D U-Net with a ResNet-18 backbone. The training loss function was Dice, validation metric was intersection over union (IoU), the gradient descent used Adam optimization with initial learning rate of 0.0001, and weights were initialized randomly. The model was trained using 20 WT datasets (40 hindpaws) with equal sex and age (from 2-6-months) distribution, each split into 6 subvolumes (3 per hindpaw) of 200 x 200 x 200 voxels with a 25% randomized validation set of subvolumes (30 validation, 120 total) to avoid overfitting. These 3D tiles were positioned evenly on the tarsals, distal phalanges, and background regions. The training patch size was set at 96 x 96 x 96 voxels (Supplementary Figure 1). The model was trained over 500 epochs, taking approximately 6 hours on a Nvidia RTX 8000 GPU. Ground truth joint regions were obtained by an automatic recipe from ground truth labels, which expanded label interfaces with 3D dilation size 5 for both thickness and extent.

### 2.4 Segmentation Method Testing and Quantification

The segmentation method was tested in Amira (v2023.1.1) through generation of a recipe that incorporated the DL joint prediction (Python environment: deep-learning-environment-2022_2; architecture, weights, and custom processing files provided in Supplementary Files; tiling width, height, depth = 352 pixels without tiling overlap) with the downstream image processing recipe on intensity range 2500 – 20000 Hounsfield units. Recipe generation allowed for batch processing (<u>Apply a Recipe on a Batch of Files</u>) of the original micro-CT datasets (.am file format as image stack saved after import of initial .dcm files into Amira). The initial WT test datasets included 44 additional hindpaws that were not involved in training or validation. Computer hardware included 16 cores from an Intel® Xeon® Gold 5218 central processing unit (CPU) (Intel, Santa Clara, CA) at 2.30GHz, 128GB of double data rate fourth generation (DDR4) error correction code (ECC) random-access memory (RAM) at 2666 megatransfers (MT)/s, and 24GB of virtual graphics processing unit (vGPU/VRAM) from NVIDIA A40 (NVIDIA, Santa Clara, CA) on a 64-bit operating system running Windows 10 (Microsoft Corporation, Redmond, WA; operating system build: 19044.4780). Each hindpaw dataset (2 hindpaws) was segmented in approximately 32.7±8.42 minutes (mean ± standard deviation) without user intervention. Forepaw datasets (2 forepaws) were segmented in approximately 53.4±23.6 minutes without user intervention, where increased segmentation time can be attributed to additional structures within the original imaging datasets (i.e., spine and ribs), which are not present in the more distally isolated hindpaws and inflates the segmentation time in the absence of preceding volume editing steps.

Quantification of accuracy was performed by visual inspection (HMK) to identify correct segmentation or error type based on expected bone anatomy (Hindpaw: (Bab et al., 2007a; Kenney et al., 2022b); Forepaw: (Bab et al., 2007b)). Accuracy was calculated as a percentage by:

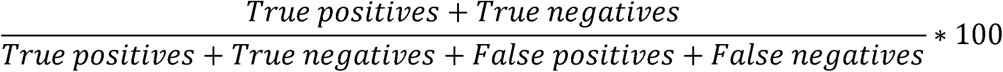

where true positives were correctly segmented bones, true negatives equal 0 (there are no circumstances in which bones should be missing and background was not relevant for quantification), false positives were over-split bones, and false negatives were over-connected bones. Accuracy was determined to be an appropriate quantitative metric given the single class problem identifying spaces between bones and true negatives were non-contributory to the outcomes thus with lower risk of overestimating performance. The automated segmentation method does not include bone naming, the bone names are later associated with segmented materials manually by the user.

Evaluation of the hindpaws involved in this study have confirmed the fixed fusion in the tarsals of the navicular and lateral cuneiform (NAVLAT) in C57BL/6 mice (Richbourg et al., 2017), but also determined that the adjacent intermediate cuneiform (INT) may also be variably fused with the NAVLAT structure (NAVLATINT) (Bab et al., 2007a; Kenney et al., 2022b). Similar variable fusion was appreciated in the carpal region of forepaws where the trapezoid (ZOID; lesser multangular) and centrale (CENT) bones may present as either a single fused structure (CENTZOID) or subdivided into their individual bones in particular forepaws. Additional carpal bones investigated for segmentation accuracy included the trapezium (ZIUM; greater multangular), capitate (CAP), hamate (HAM), triquetrum (TRI; triangular), pisiform (PIS), scaphoid (navicular) / lunate (SCAPHATE; fixed fusion), and falciformis (FALC). The forepaw metacarpals (MET-F; 1-5), proximal phalanges (PP-F; 1-5), distal phalanges (DP-F; 2-5), and sesamoids (S-F; 1-10) are numbered lateral to medial, as opposed to hindpaw counterparts (metatarsals (MET-H), PP-H, DP-H, and S-H) which are numbered medial to lateral (Kenney et al., 2022b). Along with NAVLATINT, additional tarsal bones were evaluated for hindpaws, as previously described (Kenney et al., 2022b; Kenney et al., 2024b), including the calcaneus (CALC), cuboid (CUB), medial cuneiform (MED), talus (TAL), and tibiale (TIB). Note the overall cohort accuracy quantifications vary slightly when comparing assessment of average accuracy per dataset with variable number of bones due to anatomical fusions versus accuracy calculated based on total individual bones analyzed.

### 2.5 Statistical Analysis

Statistical analysis including 3-way or 2-way mixed-effects analysis with interaction effects or Sidak’s multiple comparisons and Fisher’s exact test were performed as appropriate in GraphPad Prism (v10.2.0; San Diego, CA, USA). Males (2-8-months) and females (2-5-months) were analyzed separately given the distinct timeframes of evaluation based on early TNF-Tg female mortality (Bell et al., 2019). The sample sizes of the WT hindpaws used for training/validation and methodologic testing are provided in Supplementary Table 1, along with sample size details for tested WT and TNF-Tg hindpaws and forepaws in Supplementary Tables 2-4. As certain timepoints for WT hindpaw testing incorporated accuracy evaluation for <3 hindpaws, interaction effects were reported without post-hoc multiple comparisons in analyses that included WT hindpaws. Entire or portions of hindpaws were omitted from analysis if there were imaging errors with incomplete capture of the paw, considerable motion artifact rendering the scans uninterpretable, and/or if the animal died prior to a scheduled imaging session as all data was collected *in vivo*.

## 3. Results

### 3.1 Implementation of automated joint space identification improves bone segmentation accuracy

Given the heterogeneity of bone shape and architecture in complex structures such as the murine hindpaw, we build upon our systematic image processing algorithm (Kenney et al., 2022b) through DL training predictions (blue) coupled with image processing steps for robust identification of inter-bone joint spaces in micro-CT datasets (Fig. 1A-B; process described in Methods and shown in Supplementary Fig. 1). The identification of spaces between bones enabled precise bone separation and segmentation of individual hindpaw bones (separate colors) for downstream analysis (Fig. 1C). For the added DL facilitation component (image processing algorithms + DL), the training and validation datasets (WT) consisted of equal ages (2-6 months of age, n=8 hindpaws per age) and sex (n=20 hindpaws per sex). The remainder of the WT hindpaws (n=44, from 2-8 months of age, excluding 6 months as all used for training and validation) served as the test datasets to quantify accuracy of bone segmentation (Fig. 1D).

**Figure 1.**
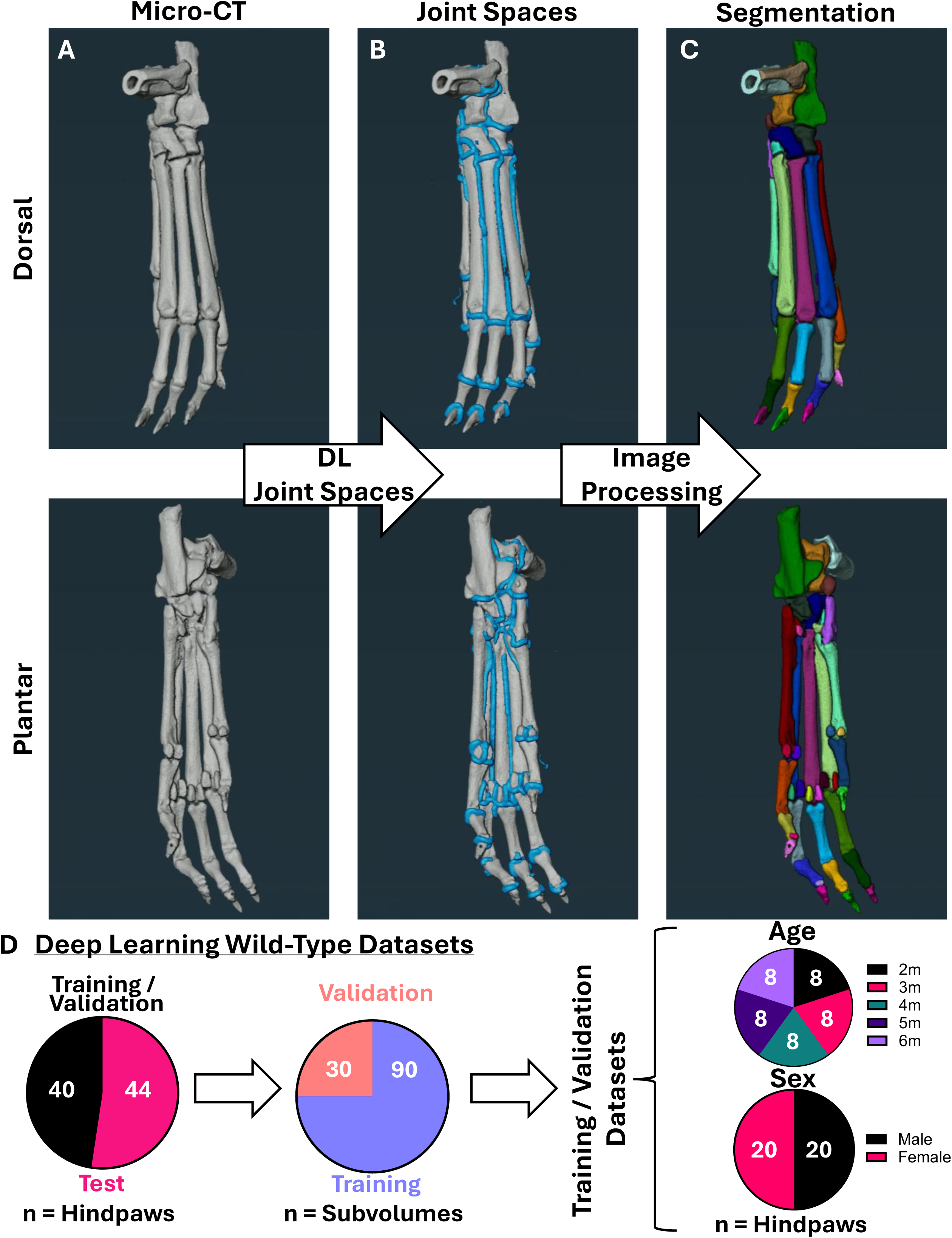
Automated joint space detection via strategic image processing and deep learning predictions for bone segmentation. Murine micro-CT datasets with visualization from the dorsal (top) and plantar (bottom) surfaces **(A)** were processed for subsequent automated identification of joint spaces (blue) **(B)** using a DL model (described in Supplementary Figure 1) developed from gold-standard bone segmentations (Kenney et al., 2022b; Kenney et al., 2024b). Final successful bone separation (bone-specific colors) was accomplished through an additional combination of image processing steps, including black top-hat (Kenney et al., 2022b), structure enhancement (Frangi et al., 1998), and tensor voting (Martinez-Sanchez et al., 2014) for robust joint space identification to label individual bones **(C)**. Training and validation (n=40 hindpaws) of the DL component were performed with WT murine hindpaws of equal age (from 2-6-months, n=8 hindpaws each timepoint) and sex (n=20 hindpaws male/female) distribution with a randomized 25% of subvolumes used for validation (3 subvolumes per hindpaw, total 120 subvolumes). The remaining WT hindpaws (n=44) were evaluated as test cases for further analysis **(D)**. The combination of the DL model and image processing algorithms were evaluated using previously published and publicly available datasets (Kenney et al., 2024a; Kenney et al., 2024b).

Along with implementation in WT hindpaws, we also tested the automated segmentation approach on hindpaws from TNF-Tg mice with spontaneous inflammatory-erosive arthritis. The novel segmentation algorithm automatically detected the joint spaces (blue, left) for individual bone separation (colors, right) across both sexes and genotypes (Fig. 2A-D). For segmentation accuracy of individual bones shown in Supplementary Tables 5 and 6, WT outperformed TNF-Tg datasets for both males (WT 98.4% vs TNF-Tg 93.1%, *p<0.0001*) and females (WT 98.7% vs TNF-Tg 92.1%, *p<0.0001*). The source of error was demonstrated visually as incomplete closure of joint spaces (arrows in white dashed box), thus inadvertently over-connecting 2 distinct bones into a single segmentation (Fig. 2C-D) and may represent sequelae of chronic damage leading to joint fusion where the space between bones no longer exists. In fact, the difference in accuracy between WT and TNF-Tg datasets becomes more pronounced across time as the arthritic severity increases (Fig. 2E-F), especially in the tarsal bones (Fig. 2G-H, yellow = increased accuracy, green = decreased accuracy) that typically serve as reliable biomarkers for the progression of bone erosion (Kenney et al., 2024b). However, compared to our prior SA segmentation approach, there was remarkable improvement in dataset accuracy overall (Fig. 2E-F; WT male: SA 79.39±5.73% vs DL 98.16±1.47%, *p<0.0001*; WT female: SA 79.16±4.84% vs DL 99.19±1.63%, *p<0.0001*), demonstrating the robust methodologic advancements both in automaticity and reliability.

**Figure 2.**
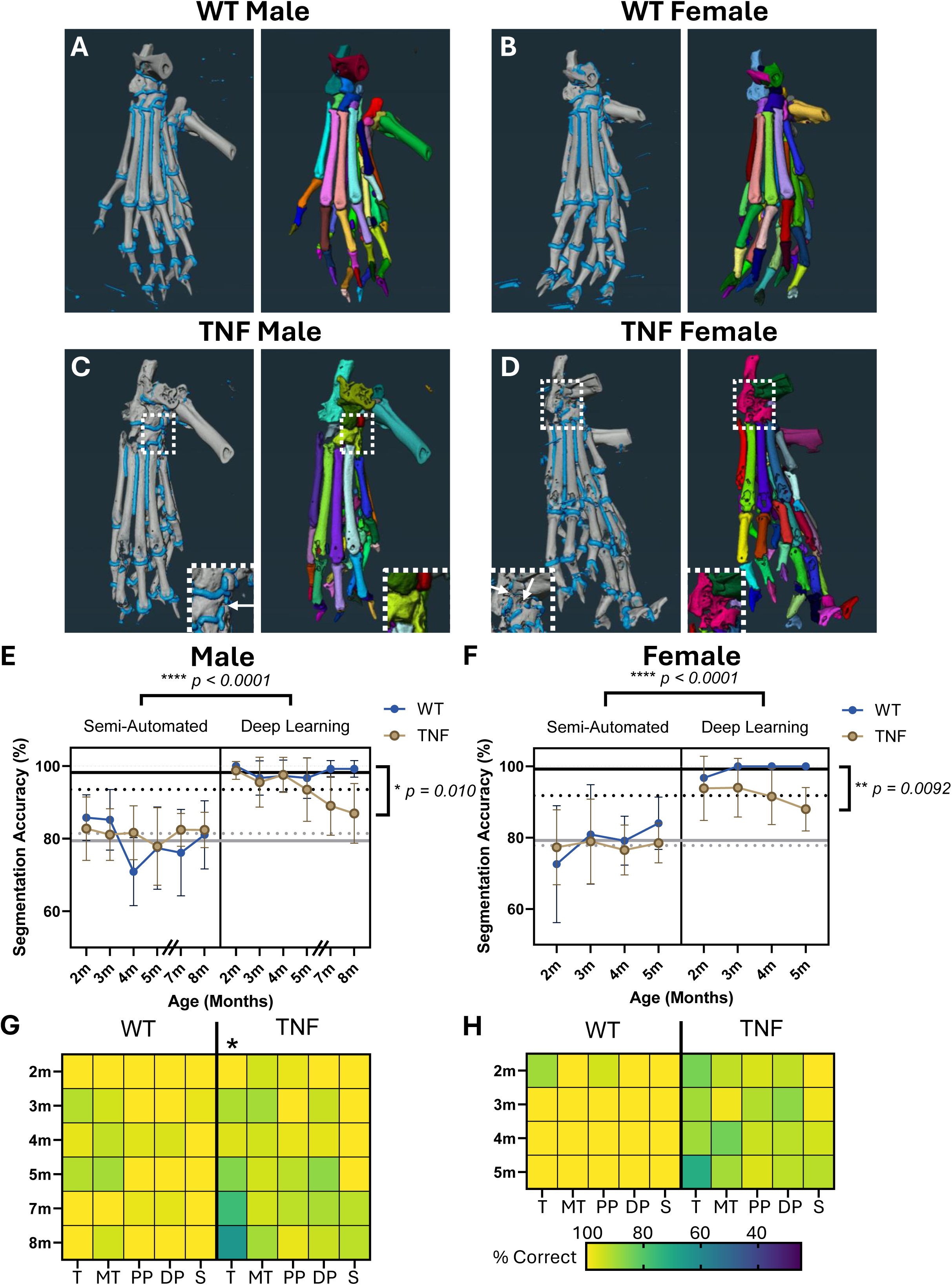
Implementation of automated joint space identification with deep learning facilitation improves bone segmentation accuracy. Following development of the automated joint space detection, we applied the DL model (left: blue joint spaces; right: bone-specific segmentation colors) to the remaining test cases for WT males and females **(A-B)**. We also assessed performance on age-matched cohorts (males: 2-8-months; female: 2-5-months) of TNF-Tg mice **(C-D)** with progressive inflammatory-erosive arthritis associated with early onset mortality in females (Bell et al., 2019). Note the 6-month male timepoint was omitted as all WT datasets were utilized for training and validation so were not included in the DL testing cohort. Compared to our previous SA segmentation algorithms (Kenney et al., 2022b; Kenney et al., 2024b), the segmentation accuracy (bones correctly segmented / total bones) was remarkably improved for both WT and TNF-Tg datasets with the DL approach regardless of sex (average accuracy lines: solid black = DL WT, dashed black = DL TNF-Tg, solid grey = SA WT, dashed grey = SA TNF-Tg). However, the accuracy of TNF-Tg segmentations notably declined with time and associated progressive joint damage compared to WT, although continued to outperform the SA method **(E-F)**. Heatmaps of accuracy specified to bone compartments (T = tarsals, MT = metatarsals, PP = proximal phalanges, DP = distal phalanges, S = sesamoids) demonstrate the increased error rate in TNF-Tg mice is predominately localized to the tarsal region (light = high, dark = low accuracy) **(G-H)**. Inset images **(C-D)** highlight the source of error with disconnected joint spaces (arrows, left image) leading to over-connected bones (colors, right image). In fact, the errors were predominately over-connected (2+ bones segmented as 1 material; noted in Supplementary Figure 2), which may represent the pathologic process of joint fusions with increasing arthritic severity. Statistics: 3-way mixed-effects analysis (SA vs DL; method x genotype x time) **(E-F)**, 2-way mixed-effects analysis (WT vs TNF; genotype x time) **(E-H)**; *\*\*\*\*p<0.0001, **p<0.01, *p<0.05*.

### 3.2 Flexible application of the segmentation method to forepaws highlights pronounced joint destruction and bone fusions in TNF-Tg mice with rapid reduction in segmentation accuracy

We further extended the application of the novel segmentation method to murine forepaws with unique bone size and anatomy. For orientation, we provide a model WT forepaw with each individual bone separated by color and bone-specific nomenclature indicated from different viewpoints (Fig. 3). Prior investigation in TNF-Tg mice has primarily focused on the hindpaw, while here we demonstrate the architecture of murine forepaws in both WT and TNF-Tg mice, particularly highlighting the carpals (yellow dashed circle) and sesamoids (blue dashed circle) that exhibit visually profound erosive disease, particularly in TNF-Tg females (Fig. 4A-D). As such, comparison of hindpaw and forepaw segmentation accuracy showed marked reduction in forepaws (paw type effect *p<0.0001*) primarily driven by steep decline with increased age and disease severity in TNF-Tg datasets (Fig. 4E-F; paw x genotype effect *p=0.0083*; male forepaws: WT 87.29±2.07% vs TNF-Tg 72.65±11.70%, *p<0.0001*). Similar to hindpaws, the decline in TNF-Tg segmentation accuracy with aging and disease severity is more pronounced in carpals, along with the sesamoids (Fig. 4G-H, Supplementary Tables 7 and 8) that may be driven by enhanced erosive activity at the adjacent articulation of the MET-F and PP-F (metacarpophalangeal joint). Evaluation of error type revealed that TNF-Tg forepaws tend to exhibit a higher proportion of completely eroded bones compared to hindpaws (Supplementary Fig. 2, red “missing”). The severe erosions in TNF-Tg forepaws are further demonstrated by representative images across time that highlight the carpal region (white arrows) and the progressive complete dislocation of the paw from the forearm (yellow arrows) most notable in TNF-Tg females (Supplementary Fig. 3).

**Figure 3.**
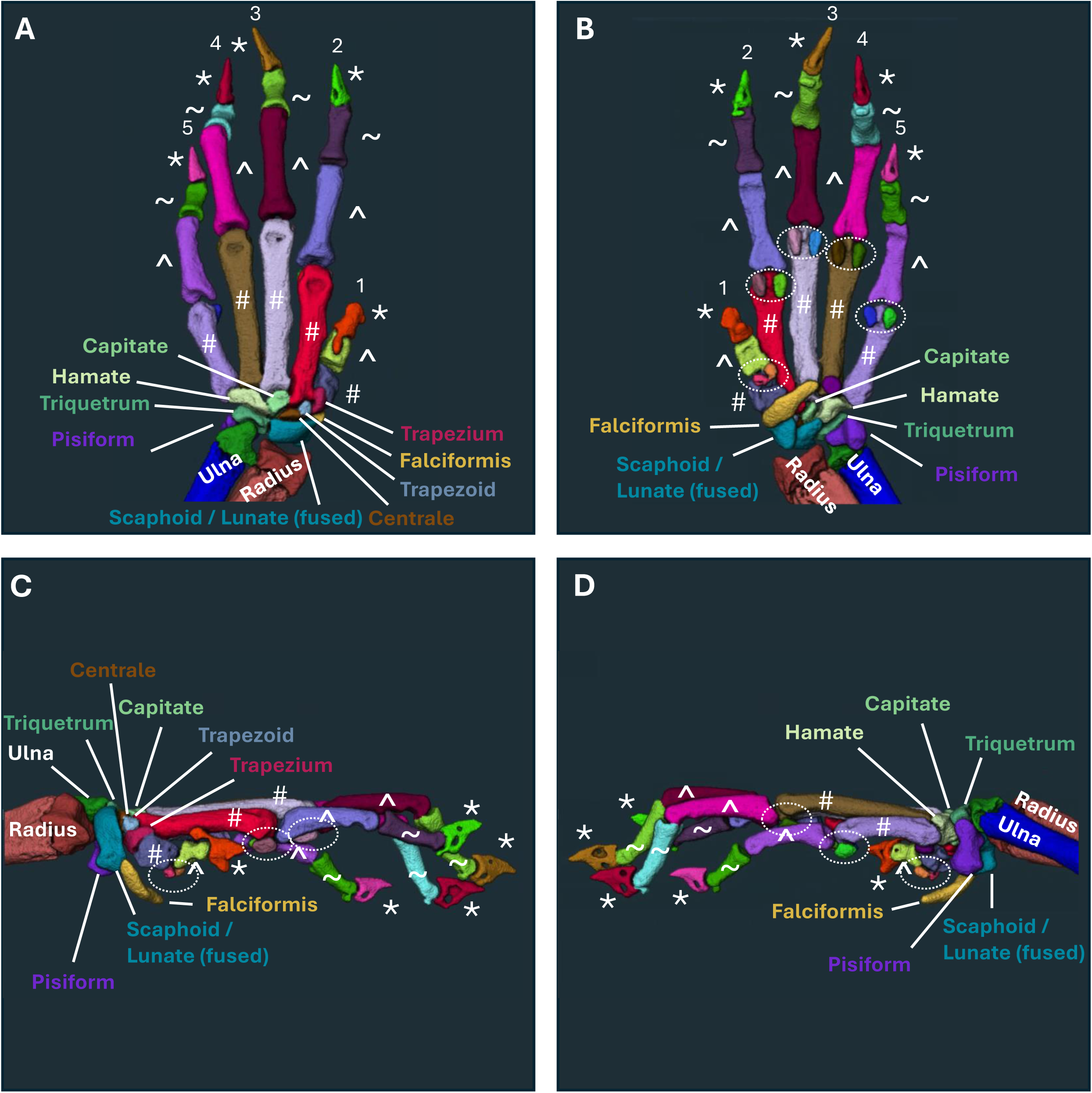
Flexible application of joint space deep learning segmentation to other complex structures highlights murine forepaw bone anatomy. To evaluate the potential for the joint space segmentation DL model to automatically separate bones in additional complex structures beyond the hindpaw, we evaluated the utilization in corresponding forepaw micro-CT datasets visualized from the dorsal **(A)**, plantar **(B)**, lateral **(C)**, and medial **(D)** surfaces with colors representing individual segmented bones. We identified the potential for accurate segmentation of the forepaw bones, including distinct carpals, metacarpals (#), proximal phalanges (^), distal phalanges (∼), sesamoids (dashed circles), and claws (*) with bone-specific labeling corresponding with known forepaw anatomy (Bab et al., 2007b).

**Figure 4.**
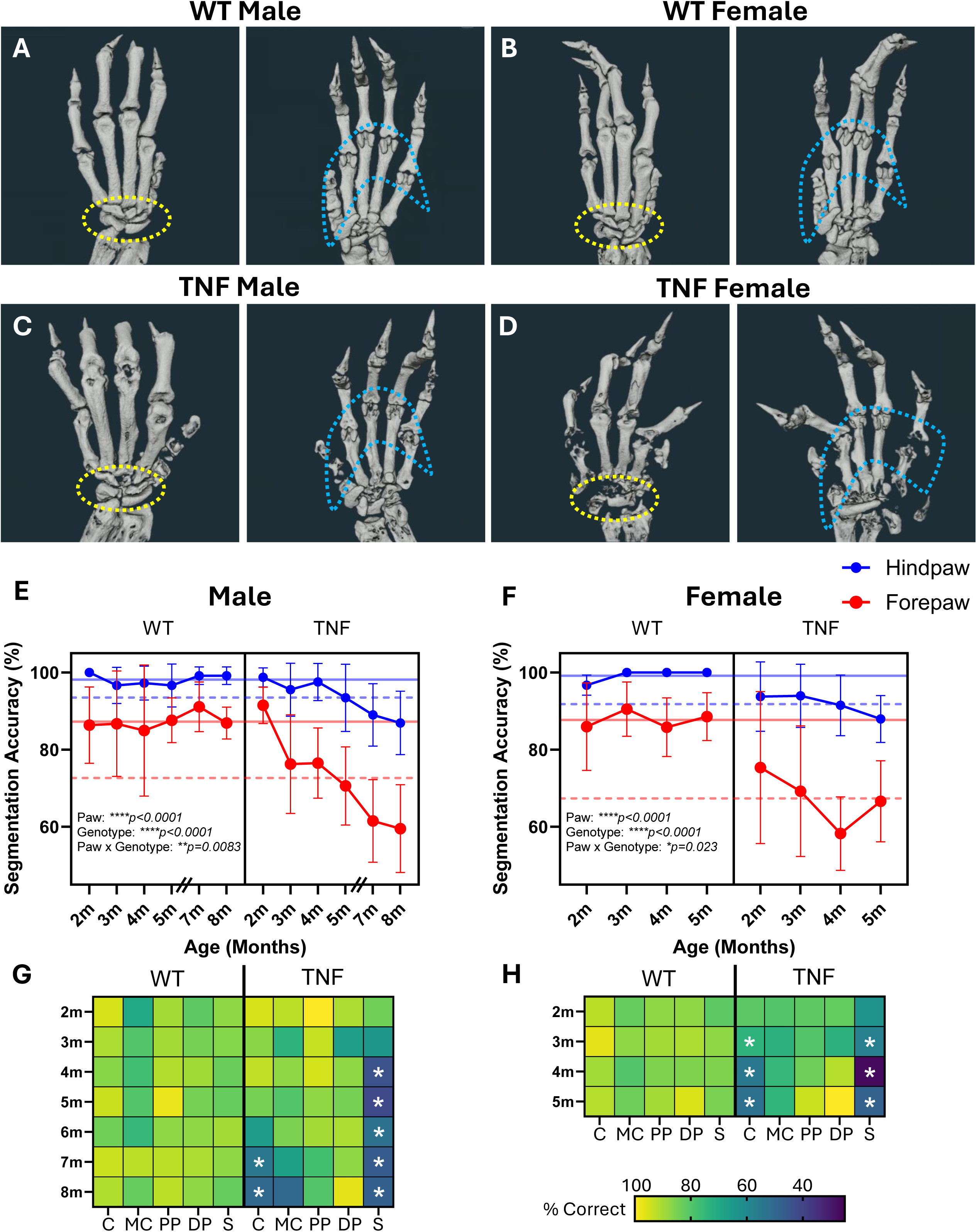
TNF-Tg mice exhibit pronounced forepaw joint destruction and bone fusions with rapid reduction in segmentation accuracy. Given the complexity and small architecture of murine forepaws highlighted by dorsal (left) and plantar (right) visualization of micro-CT images from WT male and female mice **(A-B)**, the anatomy and associated arthritis in TNF-Tg mice **(C-D)** has not been previously assessed. Bone destruction was more prominent in the murine forepaw than hindpaw where the forepaws display earlier severe bone erosions. Application of our novel joint space DL approach provided an initial opportunity to evaluate these complex structures by reducing the analytical challenges with achievement of >85% accuracy of WT forepaws, although with deficient accuracy compared to hindpaws (average accuracy lines: solid blue = WT hindpaw, dashed blue = TNF-Tg hindpaw, solid red = WT forepaw, dashed red = TNF-Tg forepaw). In addition, TNF-Tg forepaws showed rapid and dramatic decline in segmentation accuracy **(E-F)** due to errors localized to the carpals (yellow dotted circles in **A-D**) and sesamoids (blue dotted circles in **A-D**) over time with increasing severity of joint disease as shown by heatmaps (light = high, dark = low accuracy) of bone compartments (C = carpals, MC = metacarpals, PP = proximal phalanges, DP = distal phalanges, S = sesamoids) **(G-H)**. Note the 6-month male timepoint was omitted in **E** as all WT hindpaw datasets were utilized for training and validation so were not included in the DL testing cohort. Statistics: 3-way mixed effects analysis (hindpaw vs forepaw, WT vs TNF; paw type x genotype x time) **(E-F)**, 2-way mixed effects analysis with Sidak’s multiple comparisons (WT vs TNF; genotype x time) **(G-H)**; *\*\*\*\*p<0.0001, **p<0.01, *p<0.05*.

## 4. Discussion

To the end of fully automated analyses of bone volumes in mice, we generated further improvements in segmentation of micro-CT data in complex structures, particularly murine hindpaws, for downstream analytics. The strategy was to target the articulating joint spaces to create boundaries for bone separation, where focusing on the negative space between bones enabled flexible implementation in alternative structures like the forepaw, since the approach was not particular to the shape and anatomy of the distinct hindpaw bones. Although the segmentation accuracy decreased when performed on forepaws, especially with severe erosion, WT datasets still demonstrated bone accuracy of >85%. Well-described corrective processes (Kenney et al., 2022b) could be applied to create forepaw model datasets for DL training, dramatically lowering barriers for creation of structure-specific algorithms. This novel approach also allowed for application to TNF-Tg paws with severe and progressive inflammatory-erosive arthritis. In TNF-Tg paws, the decline in segmentation accuracy was striking over time, corresponding to the progressive increase in bone erosions and eventual pathologic bone-bone fusions from remodeling eroded surfaces with increased age. Thus, the remarkably successful application of an automated and highly accurate segmentation model in WT structures has the potential to guide future applications in disease models or other complex joints. Further investigation will focus on optimizing segmentation of arthritic joints that may quantify pathologic effects of bone erosions and fusions to identify disease biomarkers.

Despite the successful utilization of micro-CT imaging to monitor erosion of small bones in pre-clinical arthritis models (Cambre et al., 2018; Kenney et al., 2022b; Kenney et al., 2023; Kenney et al., 2024b; Proulx et al., 2007), there has been limited application of CT modalities in clinical evaluation. In particular for rheumatoid arthritis, scoring systems are primarily implemented for MRI (Dakkak et al., 2020), ultrasound (De Miguel et al., 2017; Dimanti et al., 2018), and/or conventional X-ray (Ornbjerg and Ostergaard, 2019) to generate semi-quantitative and user-dependent measures of disease severity, often in conjunction with clinical metrics (England et al., 2019). As CT is considered the gold standard reference for evaluation of bone integrity (Dohn et al., 2006; Dohn et al., 2008), further optimization of clinically translatable analytical approaches promises to provide tremendous benefit for reliable and longitudinal quantitative assessment of bone volumes both to inform measures of disease severity and to evaluate treatment response. Although imaging modalities such as MRI provide a greater breadth of information, including regions of inflammation, bone marrow changes, and soft tissue pathology, novel CT imaging approaches with multi-energy inputs (Jans et al., 2018) provides promise for extending CT utilization beyond the bone architecture. Similar to our recent identification of bone-specific biomarkers in pre-clinical arthritis models (Kenney et al., 2024b), a detailed clinical effort investigating purely quantitative metrics of bone erosion would be a major advancement in disease monitoring.

While our current work provides a foundation for clinical implementation, given the potential for flexible application to novel structures by targeting joint spaces, a primary limitation is the reliance on a well-documented pre-clinical, research-oriented software in Amira not intended for clinical diagnosis. However, the underlying algorithms and strategic design can be readily implemented in alternative software environments through the detailed Methods provided. Regardless of the research software used, incorporation into clinical use (rather than investigation) requires translation efforts that meet the regulatory requirements for introduction to clinical practice. For the application of the novel segmentation strategy, it is also important to consider the potential limitations in differential image resolution, where we have previously described that image resolution (i.e., voxel / structure size) is a key determinant of segmentation accuracy using solely image processing algorithms (Kenney et al., 2022b). In fact, this is possibly associated with the slight reduction in segmentation accuracy of forepaws, where the decreased size of the forepaw structures would inherently produce relatively reduced image resolution compared to hindpaws. Lastly, expanding the described methods beyond a single-class bone separation approach to a more robust multi-class analytical tool that includes predicted bone names based on structural architecture or coordinate location in fixed anatomy (i.e., akin to an atlas tree) will certainly provide essential improvements and likely enhance method adoption.

In conclusion, we have designed a novel image processing and DL facilitated micro-CT segmentation strategy to isolate individual bones within complex structures. This innovation demonstrates a remarkable improvement in both automaticity and segmentation accuracy compared to our recently created SA workflow (Kenney et al., 2022b), that served herein as the foundation for production of numerous gold-standard segmentations to train DL models and optimize the current improvements. Similarly, the enhanced segmentation methods provide an opportunity for application to distinct structures even in different species. We further urge the incorporation of such strategies into clinical research as it promises eventual benefits for patient care.

## Supporting information

Amira Recipes and DL Files

## 6. Additional Information

### 6.1 Data Availability

As described in section *2.2* *Micro-CT Image Collection*, the hindpaw data was previously published (Kenney et al., 2022b; Kenney et al., 2023; Kenney et al., 2024b) and publicly available (Kenney et al., 2024a). For the purposes of the described study, the corresponding forepaw data has also been made publicly available at the Zenodo repository (Kenney et al., 2025).

### 6.2 Author Contributions

HMK – conceptualization, methodology, formal analysis, investigation, data curation, visualization, writing original draft; DL – methodology, software, validation, visualization, writing reviewing & editing; RB – methodology, software, validation, visualization, writing reviewing & editing; LS – resources, data curation, writing reviewing & editing; RDB – supervision, writing reviewing & editing; CTR – supervision, writing reviewing & editing; EMS – resources, supervision, funding acquisition, writing reviewing & editing; HAA – resources, supervision, writing reviewing & editing; RWW – conceptualization, methodology, supervision, writing reviewing & editing

## 6.4 Acknowledgements

We would like to thank the faculty and staff of the Histology, Biochemistry, and Molecular Imaging core, the Biomechanics, Biomaterials, and Multimodal Tissue Imaging core, and the Center for Musculoskeletal Research at the University of Rochester Medical Center for their contributions to this work.

