## Supplementary Figures for "Automated Joint Space Detection Improves Bone Segmentation Accuracy"

### Slide 1
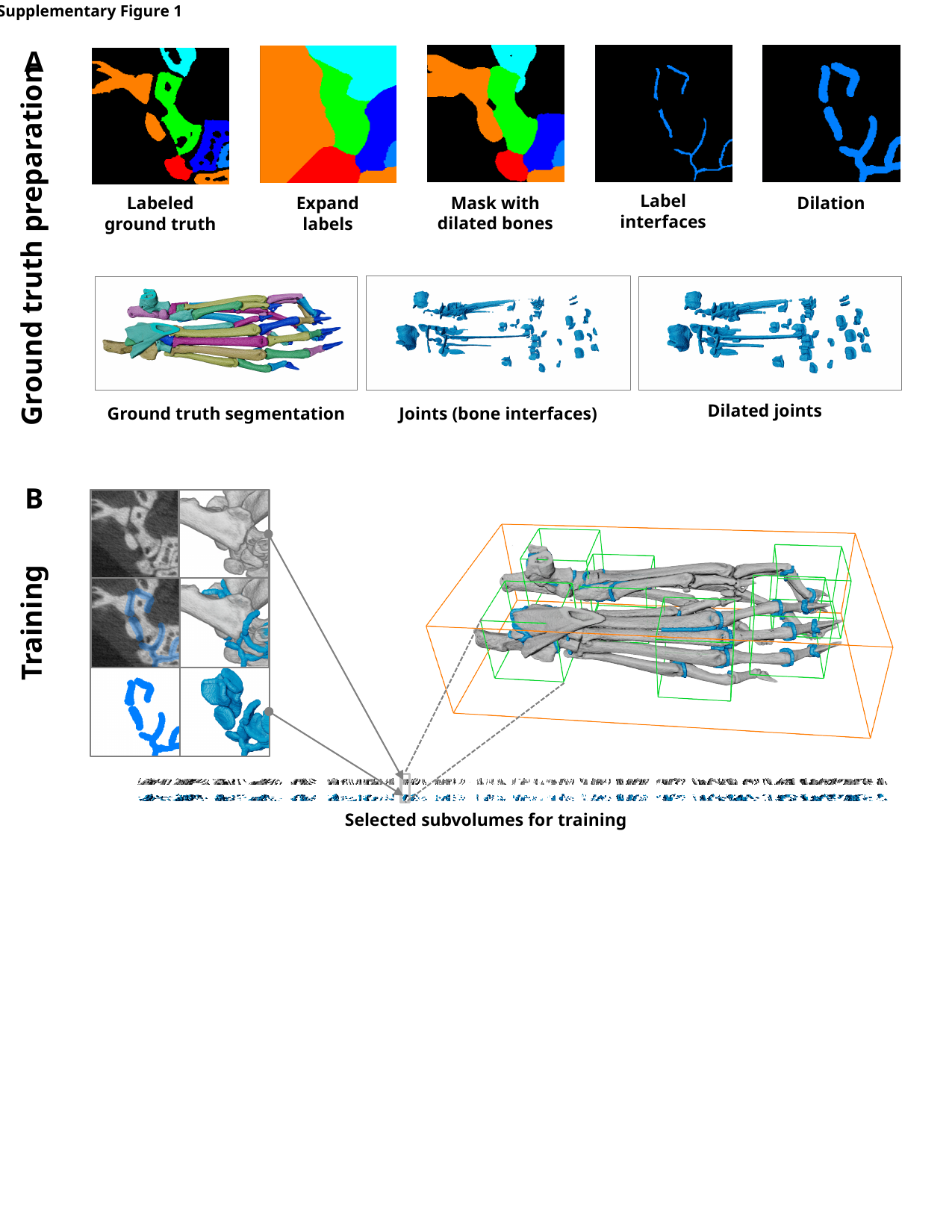

Supplementary Figure 1
A
Label interfaces
Mask with dilated bones
Dilation
Expand labels
Labeled ground truth
Ground truth preparation
Dilated joints
Ground truth segmentation
Joints (bone interfaces)
B
Training
Selected subvolumes for training

### Slide 2
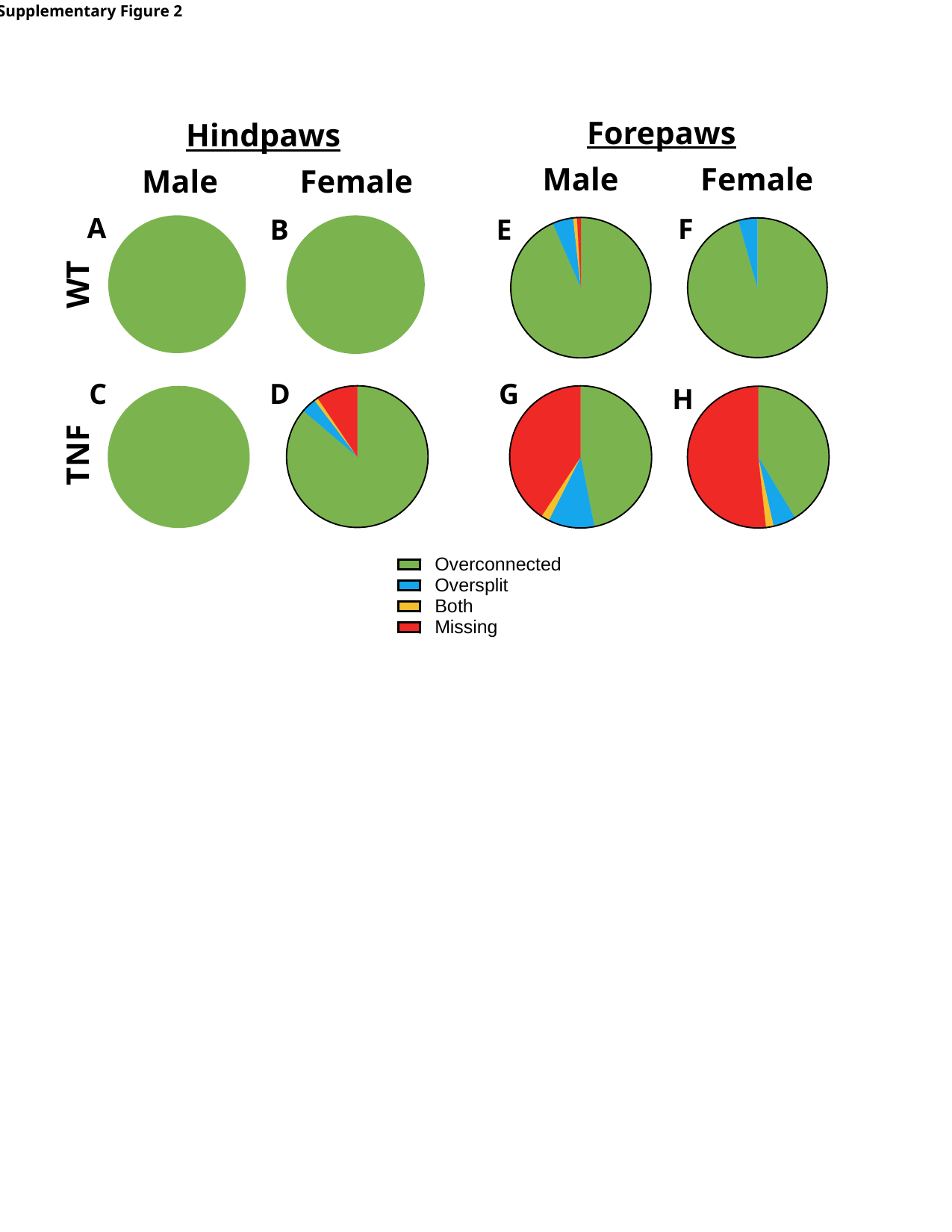

Supplementary Figure 2
Forepaws
Hindpaws
Male
Female
Male
Female
A
F
B
E
WT
G
C
D
H
TNF

### Slide 3
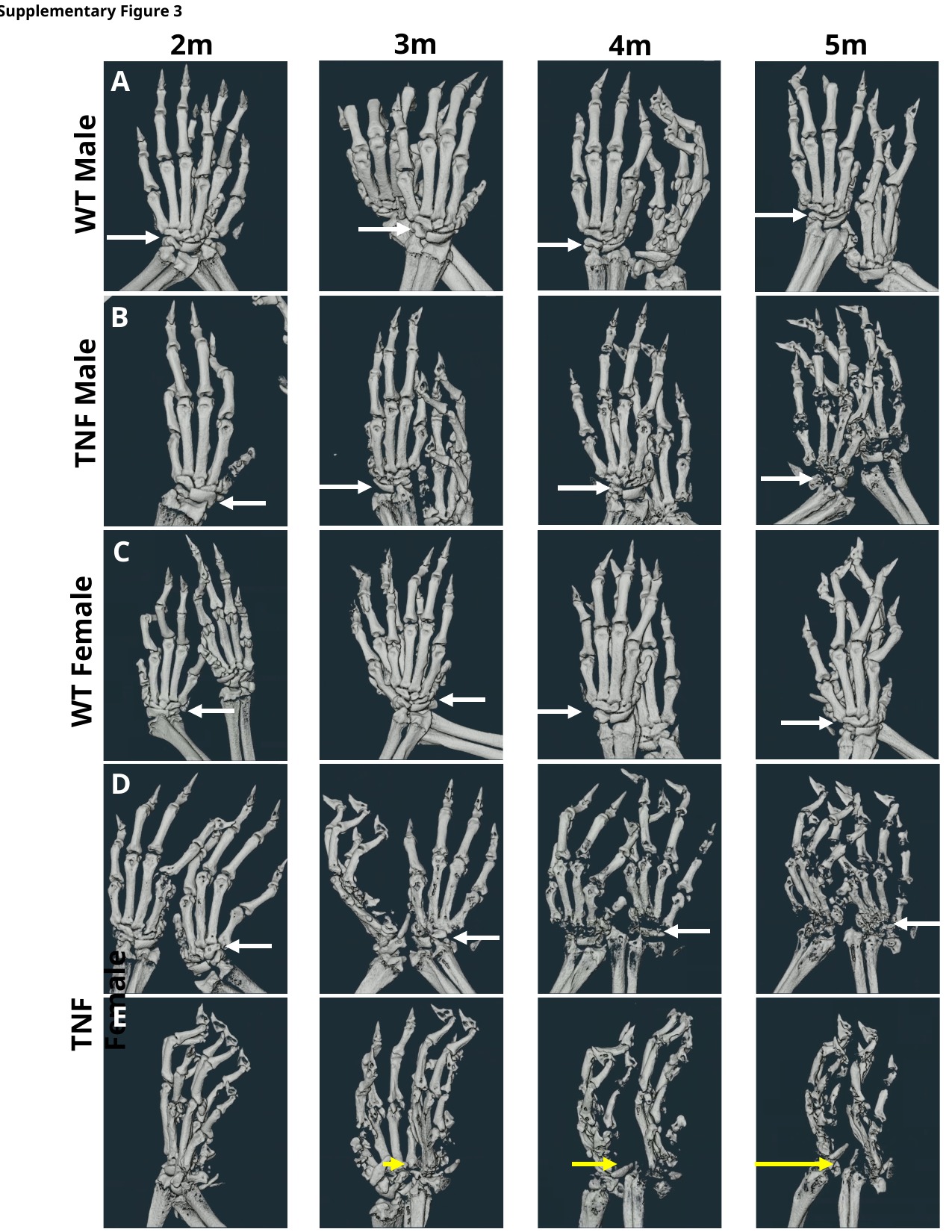

Supplementary Figure 3
3m
2m
5m
4m
A
WT Male
B
TNF Male
C
WT Female
D
TNF Female
E
