## Supplementary Figure Legends for "Automated Joint Space Detection Improves Bone Segmentation Accuracy"

**Supplementary Figure 1. Development and training of joint detection deep learning model.** Ground truth joint regions were obtained from initial ground truth bone segmentations by an automatic recipe using Amira, which combines label expansion, extraction of label interfaces, masking, and dilation **(A)**. For each of the 20 training micro-CT datasets (40 hindpaws), 6 subvolumes of 200 x 200 x 200 voxels were manually extracted from tarsals, distal phalanges, and background regions, evenly split between left and right paws (3 patches per hindpaw). The resulting 120 subvolumes were then used as input for a 3D segmentation Amira training module along with corresponding labeled joint regions as ground truth target. A randomized subset of 25% patches was used for validation to control model overfitting during the training **(B)**.

**Supplementary Figure 2. Distinct distribution of error types between hindpaws and forepaws.** Similar to the SA segmentation algorithm previously developed (Kenney et al., 2022; Kenney et al., 2024), the joint space DL model produced the greatest proportion of errors by over-connecting bones (green, 2+ bones segmented as 1 material) most notable in the hindpaws **(A-D)** or WT forepaws **(E-F)**. As noted in Figure 2, overconnected errors will occur if there is a gap in the detected joint space that may occur for various reasons, including greater bone proximity than image resolution, motion artifact blurring the joint space, or bone remodeling in the context of arthritis leading to joint fusions. Interestingly, TNF-Tg forepaws **(G-H)** exhibit a remarkably increased proportion of “missing” bones (red), meaning the bone was completely absent from the segmentation, which is likely attributed to a combination of severe erosions and deficiencies in image resolution given the relatively decreased size of forepaw bones, especially carpals and sesamoids as predominant source of error (Figure 4), compared to those of hindpaws. Additional types of errors include over-split (blue, 1 bone segmented as 2+ materials) or “both” overconnected and oversplit (orange). Pie charts represent proportions of total errors attributed to specific error subtypes.

**Supplementary Figure 3. Evaluation of progressive TNF-Tg forepaw arthritis with severe bone erosions and joint dislocations.** To visualize the structural changes in forepaws over time, we provided representative images of the dorsal surface from WT male **(A)**, TNF-Tg male **(B)**, WT female **(C)**, and TNF-Tg female **(D)** forepaws over time from 2-5-months (left to right) to particularly highlight the carpal region (white arrows). Note the severe bone erosions and remodeling that occurs by approximately 4-months in females and 5-months in males. These time periods predate the typical onset of severe bone erosions in hindpaws at approximately 5-months in females and 7-8-months in males (Kenney et al., 2024). A side view of TNF-Tg female forepaws is also shown to demonstrate the progressive dislocation of the entire paw from the forearm (yellow arrows) associated with the joint destruction **(E)**.
