## Supplementary Tables for "Automated Joint Space Detection Improves Bone Segmentation Accuracy"

**Supplementary Tables and Legends**

|  | **WT Male** | | | **WT Female** | | |
| --- | --- | --- | --- | --- | --- | --- |
| **Age (Months)** | **Training/**  **Validation**  **(Hindpaws)** | **Testing**  **(Hindpaws)** | **Omitted**  **(Hindpaws)** | **Training/**  **Validation**  **(Hindpaws)** | **Testing**  **(Hindpaws)** | **Omitted**  **(Hindpaws)** |
| 2 | 4 | 2 | 2 | 4 | 4 | 0 |
| 3 | 4 | 4 | 0 | 4 | 2 | 2 |
| 4 | 2 | 6 | 0 | 6 | 2 | 0 |
| 5 | 2 | 6 | 0 | 6 | 2 | 0 |
| 6 | 8 | 0 | 0 |  |  |  |
| 7 | 0 | 8 | 0 |  |  |  |
| 8 | 0 | 8 | 0 |  |  |  |
| **Total** | **20** | **34** | **2** | **20** | **10** | **2** |

**Supplementary Table 1. Sample sizes of WT hindpaws for DL training, validation, and methodologic testing.** Sample sizes in number of hindpaws are provided across age (months 2-8) and organized by datasets used for DL training/validation, total methodologic testing, or those omitted either due to imaging error, severe motion artifact, and/or death prior to scheduled micro-CT scan. Black cells from months 6-8 for females indicate the planned termination of scans after 5-months due to early mortality of TNF-Tg experimental counterparts.

|  | **TNF Male** | | **TNF Female** | |
| --- | --- | --- | --- | --- |
| **Age (Months)** | **Testing**  **(Hindpaws)** | **Omitted**  **(Hindpaws)** | **Testing**  **(Hindpaws)** | **Omitted**  **(Hindpaws)** |
| 2 | 8 | 0 | 14 | 0 |
| 3 | 8 | 0 | 14 | 0 |
| 4 | 8 | 0 | 10 | 4 |
| 5 | 8 | 0 | 10 | 4 |
| 6 | 8 | 0 |  |  |
| 7 | 8 | 0 |  |  |
| 8 | 8 | 0 |  |  |
| **Total** | **56** | **0** | **48** | **8** |

**Supplementary Table 2. Sample sizes of TNF-Tg hindpaws for methodologic testing.** Sample sizes in number of hindpaws are provided across age (months 2-8) and organized by datasets used for total methodologic testing or those omitted either due to imaging error, severe motion artifact, and/or death prior to scheduled micro-CT scan. Black cells from months 6-8 for females indicate the planned termination of scans after 5-months due to early mortality of TNF-Tg female mice.

|  | **WT Male** | | **WT Female** | |
| --- | --- | --- | --- | --- |
| **Age (Months)** | **Testing**  **(Forepaws)** | **Omitted**  **(Forepaws)** | **Testing**  **(Forepaws)** | **Omitted**  **(Forepaws)** |
| 2 | 8 | 0 | 8 | 0 |
| 3 | 8 | 0 | 8 | 0* |
| 4 | 7 | 1 | 7 | 1 |
| 5 | 8 | 0 | 6 | 2 |
| 6 | 8 | 0 |  |  |
| 7 | 8 | 0 |  |  |
| 8 | 8 | 0 |  |  |
| **Total** | **55** | **1** | **29** | **3** |

**Supplementary Table 3. Sample sizes of WT forepaws for methodologic testing.** Sample sizes in number of forepaws are provided across age (months 2-8) and organized by datasets used for total methodologic testing or those omitted either due to imaging error, severe motion artifact, and/or death prior to scheduled micro-CT scan. Black cells from months 6-8 for females indicate the planned termination of scans after 5-months due to early mortality of TNF-Tg experimental counterparts. *At 3-months for WT females, n=1 forepaw had omitted DP-F3, PP-F3, DP-F4, and PP-F4 due to imaging error, although the remainder of the forepaw was evaluated.

|  | **TNF Male** | | **TNF Female** | |
| --- | --- | --- | --- | --- |
| **Age (Months)** | **Testing**  **(Forepaws)** | **Omitted**  **(Forepaws)** | **Testing**  **(Forepaws)** | **Omitted**  **(Forepaws)** |
| 2 | 8 | 0 | 14 | 0 |
| 3 | 6 | 2 | 14 | 0 |
| 4 | 8 | 0 | 12 | 2 |
| 5 | 8 | 0 | 10 | 4 |
| 6 | 8 | 0 |  |  |
| 7 | 8 | 0 |  |  |
| 8 | 8 | 0 |  |  |
| **Total** | **54** | **2** | **50** | **6** |

**Supplementary Table 4. Sample sizes of TNF-Tg forepaws for methodologic testing.** Sample sizes in number of forepaws are provided across age (months 2-8) and organized by datasets used for total methodologic testing or those omitted either due to imaging error, severe motion artifact, and/or death prior to scheduled micro-CT scan. Black cells from months 6-8 for females indicate the planned termination of scans after 5-months due to early mortality of TNF-Tg female mice.

|  | **WT Male** | | | | **TNF Male** | | | | **Fisher’s Exact Test** |
| --- | --- | --- | --- | --- | --- | --- | --- | --- | --- |
| **Bone** | **Correct** | **Incorrect** | **Total** | **% Correct** | **Correct** | **Incorrect** | **Total** | **% Correct** | ***p*-value** |
| **CALC** | 42 | 0 | 42 | 100 | 49 | 7 | 56 | 87.5 | *0.019 |
| **CUB** | 40 | 2 | 42 | 95.2 | 47 | 9 | 56 | 83.9 | 0.109 |
| **INT** | 19 | 1 | 20 | 95 | 18 | 10 | 28 | 64.3 | *0.016 |
| **MED** | 41 | 1 | 42 | 97.6 | 49 | 7 | 56 | 87.5 | 0.133 |
| **NAVLATINT** | 21 | 1 | 22 | 95.5 | 22 | 6 | 28 | 78.6 | 0.118 |
| **NAVLAT** | 20 | 0 | 20 | 100 | 16 | 12 | 28 | 57.1 | ***0.0005 |
| **TAL** | 40 | 2 | 42 | 95.2 | 49 | 7 | 56 | 87.5 | 0.293 |
| **TIB** | 42 | 0 | 42 | 100 | 55 | 1 | 56 | 98.2 | >0.999 |
| **MET-H1** | 40 | 2 | 42 | 95.2 | 53 | 3 | 56 | 94.6 | >0.999 |
| **MET-H2** | 40 | 2 | 42 | 95.2 | 52 | 4 | 56 | 92.9 | 0.698 |
| **MET-H3** | 41 | 1 | 42 | 97.6 | 51 | 5 | 56 | 91.1 | 0.234 |
| **MET-H4** | 40 | 2 | 42 | 95.2 | 52 | 4 | 56 | 92.9 | 0.698 |
| **MET-H5** | 40 | 2 | 42 | 95.2 | 54 | 2 | 56 | 96.4 | >0.999 |
| **PP-H1** | 42 | 0 | 42 | 100 | 53 | 3 | 56 | 94.6 | 0.258 |
| **PP-H2** | 42 | 0 | 42 | 100 | 54 | 2 | 56 | 96.4 | 0.505 |
| **PP-H3** | 42 | 0 | 42 | 100 | 53 | 3 | 56 | 94.6 | 0.258 |
| **PP-H4** | 40 | 2 | 42 | 95.2 | 54 | 2 | 56 | 96.4 | >0.999 |
| **PP-H5** | 42 | 0 | 42 | 100 | 56 | 0 | 56 | 100 | >0.999 |
| **DP-H2** | 42 | 0 | 42 | 100 | 53 | 3 | 56 | 94.6 | 0.258 |
| **DP-H3** | 42 | 0 | 42 | 100 | 49 | 7 | 56 | 87.5 | *0.019 |
| **DP-H4** | 40 | 2 | 42 | 95.2 | 53 | 3 | 56 | 94.6 | >0.999 |
| **DP-H5** | 42 | 0 | 42 | 100 | 54 | 2 | 56 | 96.4 | 0.505 |
| **S-H1** | 42 | 0 | 42 | 100 | 56 | 0 | 56 | 100 | >0.999 |
| **S-H2** | 41 | 1 | 42 | 97.6 | 56 | 0 | 56 | 100 | 0.429 |
| **S-H3** | 42 | 0 | 42 | 100 | 56 | 0 | 56 | 100 | >0.999 |
| **S-H4** | 42 | 0 | 42 | 100 | 56 | 0 | 56 | 100 | >0.999 |
| **S-H5** | 42 | 0 | 42 | 100 | 48 | 8 | 56 | 85.7 | **0.0098 |
| **S-H6** | 42 | 0 | 42 | 100 | 48 | 8 | 56 | 85.7 | **0.0098 |
| **S-H7** | 42 | 0 | 42 | 100 | 56 | 0 | 56 | 100 | >0.999 |
| **S-H8** | 42 | 0 | 42 | 100 | 56 | 0 | 56 | 100 | >0.999 |
| **S-H9** | 42 | 0 | 42 | 100 | 56 | 0 | 56 | 100 | >0.999 |
| **S-H10** | 42 | 0 | 42 | 100 | 56 | 0 | 56 | 100 | >0.999 |
| **Total** | **1259** | **21** | **1280** | **98.4** | **1590** | **118** | **1708** | **93.1** | ****<0.0001 |

**Supplementary Table 5. Individual bone accuracy of male hindpaws.** To identify the particular bones that reduce the segmentation accuracy in TNF-Tg vs WT hindpaws, details are provided on number of bones segmented correctly, incorrectly, and percent correct relative to total bones evaluated in male mice. Within the tarsal region where the primary deficits occur (Figure 2), the calcaneus (CALC), intermediate cuneiform (unfused, INT), and navicular / lateral cuneiform (unfused) demonstrated the most prominent decrease in accuracy for TNF-Tg hindpaws. Statistics: Fisher’s exact test; **p<0.05, **p<0.01, ***p<0.001, ****p<0.0001*.

|  | **WT Female** | | | | **TNF Female** | | | | **Fisher’s Exact Test** |
| --- | --- | --- | --- | --- | --- | --- | --- | --- | --- |
| **Bone** | **Correct** | **Incorrect** | **Total** | **% Correct** | **Correct** | **Incorrect** | **Total** | **% Correct** | ***p*-value** |
| **CALC** | 10 | 0 | 10 | 100 | 45 | 3 | 48 | 93.8 | >0.999 |
| **CUB** | 10 | 0 | 10 | 100 | 39 | 9 | 48 | 81.3 | 0.335 |
| **INT** | 4 | 1 | 5 | 80.0 | 26 | 8 | 34 | 76.5 | >0.999 |
| **MED** | 10 | 0 | 10 | 100 | 45 | 3 | 48 | 93.8 | >0.999 |
| **NAVLATINT** | 5 | 0 | 5 | 100 | 11 | 3 | 14 | 78.6 | 0.530 |
| **NAVLAT** | 4 | 1 | 5 | 80.0 | 21 | 13 | 34 | 61.8 | 0.637 |
| **TAL** | 9 | 1 | 10 | 90.0 | 38 | 10 | 48 | 79.2 | 0.669 |
| **TIB** | 10 | 0 | 10 | 100 | 46 | 2 | 48 | 95.8 | >0.999 |
| **MET-H1** | 10 | 0 | 10 | 100 | 47 | 1 | 48 | 97.9 | >0.999 |
| **MET-H2** | 10 | 0 | 10 | 100 | 43 | 5 | 48 | 89.6 | 0.575 |
| **MET-H3** | 10 | 0 | 10 | 100 | 42 | 6 | 48 | 87.5 | 0.577 |
| **MET-H4** | 10 | 0 | 10 | 100 | 43 | 5 | 48 | 89.6 | 0.575 |
| **MET-H5** | 10 | 0 | 10 | 100 | 46 | 2 | 48 | 95.8 | >0.999 |
| **PP-H1** | 9 | 1 | 10 | 90.0 | 46 | 2 | 48 | 95.8 | 0.440 |
| **PP-H2** | 10 | 0 | 10 | 100 | 45 | 3 | 48 | 93.8 | >0.999 |
| **PP-H3** | 10 | 0 | 10 | 100 | 40 | 8 | 48 | 83.3 | 0.328 |
| **PP-H4** | 10 | 0 | 10 | 100 | 47 | 1 | 48 | 97.9 | >0.999 |
| **PP-H5** | 10 | 0 | 10 | 100 | 48 | 0 | 48 | 100 | >0.999 |
| **DP-H2** | 10 | 0 | 10 | 100 | 45 | 3 | 48 | 93.8 | >0.999 |
| **DP-H3** | 10 | 0 | 10 | 100 | 38 | 10 | 48 | 79.2 | 0.184 |
| **DP-H4** | 10 | 0 | 10 | 100 | 46 | 2 | 48 | 95.8 | >0.999 |
| **DP-H5** | 10 | 0 | 10 | 100 | 46 | 2 | 48 | 95.8 | >0.999 |
| **S-H1** | 10 | 0 | 10 | 100 | 48 | 0 | 48 | 100 | >0.999 |
| **S-H2** | 10 | 0 | 10 | 100 | 48 | 0 | 48 | 100 | >0.999 |
| **S-H3** | 10 | 0 | 10 | 100 | 43 | 5 | 48 | 89.6 | 0.575 |
| **S-H4** | 10 | 0 | 10 | 100 | 47 | 1 | 48 | 97.9 | >0.999 |
| **S-H5** | 10 | 0 | 10 | 100 | 43 | 5 | 48 | 89.6 | 0.575 |
| **S-H6** | 10 | 0 | 10 | 100 | 46 | 2 | 48 | 95.8 | >0.999 |
| **S-H7** | 10 | 0 | 10 | 100 | 47 | 1 | 48 | 97.9 | >0.999 |
| **S-H8** | 10 | 0 | 10 | 100 | 48 | 0 | 48 | 100 | >0.999 |
| **S-H9** | 10 | 0 | 10 | 100 | 47 | 1 | 48 | 97.9 | >0.999 |
| **S-H10** | 10 | 0 | 10 | 100 | 48 | 0 | 48 | 100 | >0.999 |
| **Total** | **301** | **4** | **305** | **98.7** | **1358** | **116** | **1474** | **92.1** | ****<0.0001 |

**Supplementary Table 6. Individual bone accuracy of female hindpaws.** To identify the particular bones that reduce the segmentation accuracy in TNF-Tg vs WT hindpaws, details are provided on number of bones segmented correctly, incorrectly, and percent correct relative to total bones evaluated in female mice. Given the utilization of datasets for DL training and validation, along with the decreased timeframe to 5-months for comparison with TNF-Tg mice that exhibit early mortality (Bell et al., 2019), the total number of allocated DL testing hindpaws for WT females limits the capacity for individual bone comparisons to explain the overall decreased accuracy in TNF-Tg datasets. Statistics: Fisher’s exact test; *****p<0.0001*.

|  | **WT Male** | | | | **TNF Male** | | | | **Fisher’s Exact Test** |
| --- | --- | --- | --- | --- | --- | --- | --- | --- | --- |
| **Bone** | **Correct** | **Incorrect** | **Total** | **% Correct** | **Correct** | **Incorrect** | **Total** | **% Correct** | ***p*-value** |
| **CAP** | 49 | 6 | 55 | 89.1 | 37 | 17 | 54 | 68.5 | *0.010 |
| **CENT** | 18 | 3 | 21 | 85.7 | 11 | 15 | 26 | 42.3 | **0.0029 |
| **CENTZOID** | 26 | 8 | 34 | 76.5 | 17 | 10 | 27 | 63.0 | 0.274 |
| **FALC** | 55 | 0 | 55 | 100 | 54 | 0 | 54 | 100 | >0.999 |
| **HAM** | 46 | 9 | 55 | 83.6 | 42 | 12 | 54 | 77.8 | 0.475 |
| **PIS** | 54 | 1 | 55 | 98.2 | 54 | 0 | 54 | 100 | >0.999 |
| **SCAPHATE** | 53 | 2 | 55 | 96.4 | 41 | 13 | 54 | 75.9 | **0.0020 |
| **TRI** | 53 | 2 | 55 | 96.4 | 38 | 16 | 54 | 70.4 | ***0.0002 |
| **ZIUM** | 48 | 7 | 55 | 87.3 | 40 | 14 | 54 | 74.1 | 0.094 |
| **ZOID** | 17 | 4 | 21 | 81.0 | 9 | 17 | 26 | 34.6 | **0.0028 |
| **MET-F1** | 31 | 24 | 55 | 56.4 | 41 | 13 | 54 | 75.9 | *0.043 |
| **MET-F2** | 46 | 9 | 55 | 83.6 | 49 | 5 | 54 | 90.7 | 0.392 |
| **MET-F3** | 45 | 10 | 55 | 81.8 | 36 | 18 | 54 | 66.7 | 0.082 |
| **MET-F4** | 45 | 10 | 55 | 81.8 | 33 | 21 | 54 | 61.1 | *0.020 |
| **MET-F5** | 52 | 3 | 55 | 94.5 | 48 | 6 | 54 | 88.9 | 0.320 |
| **PP-F1** | 40 | 15 | 55 | 72.7 | 48 | 6 | 54 | 88.9 | 0.051 |
| **PP-F2** | 52 | 3 | 55 | 94.5 | 48 | 6 | 54 | 88.9 | 0.320 |
| **PP-F3** | 54 | 1 | 55 | 98.2 | 44 | 10 | 54 | 81.5 | **0.0040 |
| **PP-F4** | 53 | 2 | 55 | 96.4 | 50 | 4 | 54 | 92.6 | 0.438 |
| **PP-F5** | 53 | 2 | 55 | 96.4 | 50 | 4 | 54 | 92.6 | 0.438 |
| **DP-F2** | 44 | 11 | 55 | 80.0 | 48 | 6 | 54 | 88.9 | 0.291 |
| **DP-F3** | 47 | 8 | 55 | 85.5 | 47 | 7 | 54 | 87.0 | >0.999 |
| **DP-F4** | 46 | 9 | 55 | 83.6 | 45 | 9 | 54 | 83.3 | >0.999 |
| **DP-F5** | 53 | 2 | 55 | 96.4 | 48 | 6 | 54 | 88.9 | 0.161 |
| **S-F1** | 14 | 41 | 55 | 25.5 | 6 | 48 | 54 | 11.1 | 0.082 |
| **S-F2** | 26 | 29 | 55 | 47.3 | 6 | 48 | 54 | 11.1 | ****<0.0001 |
| **S-F3** | 54 | 1 | 55 | 98.2 | 34 | 20 | 54 | 63.0 | ****<0.0001 |
| **S-F4** | 55 | 0 | 55 | 100 | 35 | 19 | 54 | 64.8 | ****<0.0001 |
| **S-F5** | 53 | 2 | 55 | 96.4 | 35 | 19 | 54 | 64.8 | ****<0.0001 |
| **S-F6** | 53 | 2 | 55 | 96.4 | 37 | 17 | 54 | 68.5 | ***0.0001 |
| **S-F7** | 54 | 1 | 55 | 98.2 | 33 | 21 | 54 | 61.1 | ****<0.0001 |
| **S-F8** | 54 | 1 | 55 | 98.2 | 41 | 13 | 54 | 75.9 | ***0.0004 |
| **S-F9** | 53 | 2 | 55 | 96.4 | 28 | 26 | 54 | 51.9 | ****<0.0001 |
| **S-F10** | 55 | 0 | 55 | 100 | 36 | 18 | 54 | 66.7 | ****<0.0001 |
| **Total** | **1551** | **230** | **1781** | **87.1** | **1269** | **484** | **1753** | **72.4** | ****<0.0001 |

**Supplementary Table 7. Individual bone accuracy of male forepaws.** To identify the particular bones that reduce the segmentation accuracy in TNF-Tg vs WT forepaws, details are provided on number of bones segmented correctly, incorrectly, and percent correct relative to total bones evaluated in male mice. Within the carpal and sesamoid regions where the primary deficits occur (Figure 4), the capitate (CAP), triquetrum (TRI), centrale (unfused, CENT), scaphoid/lunate (SCAPHATE), trapezoid (ZOID), and sesamoids 2-10 demonstrated the most prominent decrease in accuracy for TNF-Tg hindpaws. Of note, the accuracy of sesamoids 1 and 2 is deficient for both WT and TNF-Tg datasets. Interestingly, metacarpal 1 actually showed improvements in segmentation accuracy in TNF-Tg mice, potentially due to close articulations with adjacent bones leading to overconnected errors that are mitigated with arthritic erosions. Statistics: Fisher’s exact test; **p<0.05, **p<0.01, ***p<0.001, ****p<0.0001*.

|  | **WT Female** | | | | **TNF Female** | | | | **Fisher’s Exact Test** |
| --- | --- | --- | --- | --- | --- | --- | --- | --- | --- |
| **Bone** | **Correct** | **Incorrect** | **Total** | **% Correct** | **Correct** | **Incorrect** | **Total** | **% Correct** | ***p*-value** |
| **CAP** | 28 | 1 | 29 | 96.6 | 24 | 26 | 50 | 48.0 | ****<0.0001 |
| **CENT** | 3 | 1 | 4 | 75.0 | 3 | 12 | 15 | 20.0 | 0.071 |
| **CENTZOID** | 20 | 5 | 25 | 80.0 | 19 | 16 | 35 | 54.3 | 0.056 |
| **FALC** | 29 | 0 | 29 | 100 | 50 | 0 | 50 | 100 | >0.999 |
| **HAM** | 27 | 2 | 29 | 93.1 | 29 | 21 | 50 | 58.0 | ***0.0008 |
| **PIS** | 29 | 0 | 29 | 100 | 44 | 6 | 50 | 88.0 | 0.080 |
| **SCAPHATE** | 27 | 2 | 29 | 93.1 | 41 | 9 | 50 | 82.0 | 0.312 |
| **TRI** | 29 | 0 | 29 | 100 | 29 | 21 | 50 | 58.0 | ****<0.0001 |
| **ZIUM** | 24 | 5 | 29 | 82.8 | 32 | 18 | 50 | 64.0 | 0.122 |
| **ZOID** | 3 | 1 | 4 | 75.0 | 6 | 9 | 15 | 40.0 | 0.303 |
| **MET-F1** | 20 | 9 | 29 | 69.0 | 34 | 16 | 50 | 68.0 | >0.999 |
| **MET-F2** | 26 | 3 | 29 | 89.7 | 35 | 15 | 50 | 70.0 | 0.054 |
| **MET-F3** | 26 | 3 | 29 | 89.7 | 39 | 11 | 50 | 78.0 | 0.235 |
| **MET-F4** | 25 | 4 | 29 | 86.2 | 36 | 14 | 50 | 72.0 | 0.174 |
| **MET-F5** | 24 | 5 | 29 | 82.8 | 45 | 5 | 50 | 90.0 | 0.485 |
| **PP-F1** | 17 | 12 | 29 | 58.6 | 38 | 12 | 50 | 76.0 | 0.131 |
| **PP-F2** | 27 | 2 | 29 | 93.1 | 44 | 6 | 50 | 88.0 | 0.703 |
| **PP-F3** | 26 | 2 | 28 | 92.9 | 41 | 9 | 50 | 82.0 | 0.310 |
| **PP-F4** | 27 | 1 | 28 | 96.4 | 44 | 6 | 50 | 88.0 | 0.411 |
| **PP-F5** | 29 | 0 | 29 | 100 | 43 | 7 | 50 | 86.0 | *0.043 |
| **DP-F2** | 27 | 2 | 29 | 93.1 | 41 | 9 | 50 | 82.0 | 0.312 |
| **DP-F3** | 23 | 5 | 28 | 82.1 | 43 | 7 | 50 | 86.0 | 0.747 |
| **DP-F4** | 25 | 3 | 28 | 89.3 | 46 | 4 | 50 | 92.0 | 0.697 |
| **DP-F5** | 27 | 2 | 29 | 93.1 | 41 | 9 | 50 | 82.0 | 0.312 |
| **S-F1** | 8 | 21 | 29 | 27.6 | 1 | 49 | 50 | 2.0 | **0.0011 |
| **S-F2** | 11 | 18 | 29 | 37.9 | 0 | 50 | 50 | 0 | ****<0.0001 |
| **S-F3** | 29 | 0 | 29 | 100 | 22 | 28 | 50 | 44.0 | ****<0.0001 |
| **S-F4** | 29 | 0 | 29 | 100 | 32 | 18 | 50 | 64.0 | ***0.0001 |
| **S-F5** | 29 | 0 | 29 | 100 | 35 | 15 | 50 | 70.0 | ***0.0006 |
| **S-F6** | 28 | 1 | 29 | 96.6 | 30 | 20 | 50 | 60.0 | ***0.0004 |
| **S-F7** | 29 | 0 | 29 | 100 | 35 | 15 | 50 | 70.0 | ***0.0006 |
| **S-F8** | 29 | 0 | 29 | 100 | 43 | 7 | 50 | 86.0 | *0.043 |
| **S-F9** | 29 | 0 | 29 | 100 | 21 | 29 | 50 | 42.0 | ****<0.0001 |
| **S-F10** | 25 | 4 | 29 | 86.2 | 29 | 21 | 50 | 58.0 | *0.012 |
| **Total** | **814** | **114** | **928** | **87.7** | **1095** | **520** | **1615** | **67.8** | ****<0.0001 |

**Supplementary Table 8. Individual bone accuracy of female forepaws.** To identify the particular bones that reduce the segmentation accuracy in TNF-Tg vs WT forepaws, details are provided on number of bones segmented correctly, incorrectly, and percent correct relative to total bones evaluated in female mice. Within the carpal and sesamoid regions where the primary deficits occur (Figure 4), the capitate (CAP), hamate (HAM), triquetrum (TRI), and sesamoids 1-10 demonstrated the most prominent decrease in accuracy for TNF-Tg hindpaws. Of note, the accuracy of sesamoids 1 and 2 is deficient for both WT and TNF-Tg datasets. Statistics: Fisher’s exact test; **p<0.05, ***p<0.001, ****p<0.0001*.
