## Supplementary material for "Automated Joint Space Detection Improves Bone Segmentation Accuracy": Amira Recipes and DL Files: JointsD5d5-20x6x200x3D-resnet18.py

```
import numpy as np

# These functions may be customized to specific needs, such as accommodating to
# the expectations of Neural Networks trained outside our environment

def get_model_information():
    """Model information for end user
    DO NOT MODIFY.

    Returns
    -------
        str Information message that will appear in the Deep-Learning
        models information port.
    """

    model_information = "<html>This network is a <a href=\"https://arxiv.org/abs/1505.04597\"> \
            U-Net model variant.</a></html>"
    return model_information

'''
def preprocess_input(x):
    """Preprocessing of the input array x, prior to the actual prediction by the Deep Learning Prediction module.
    This preprocessing must be the same as that which was applied during the Training of the model.

    If the normalization is not set to "None" in get_normalization_configuration(), a normalization is performed
    prior to this function call.
    Uncomment this function to use a customized preprocessing.

    Parameters
    ----------
        x : numpy array
            array to process. Shape (number, depth, width, height, channels)

    Returns
    -------
        processed array with same shape (number, depth, width, height, channels)
    """
    if not isinstance(x, np.ndarray):
        raise TypeError("'x' must be a numpy array")

    if len(x.shape) != 5:
        raise ValueError("incorrect 'x' shape, must be a 5d array with shape=(number, width, height, depth, channels)")

    # Pre-processes the data by standardizing it to zero mean and unit variance.
    # This part can be customized as the user wish.
    n = x.shape[0]
    x_tmp = x.astype(dtype=np.float32)
    x_tmp = np.reshape(x_tmp, [x_tmp.shape[0], np.product(x_tmp.shape[1:])])
    x_mean = np.mean(x_tmp, axis=1).reshape((n, 1, 1, 1, 1))
    x_std = np.std(x_tmp, axis=1).reshape((n, 1, 1, 1, 1))
    return (x-x_mean)/np.maximum(x_std, 1e-7)
'''


def postprocess_local_output(y_pred):
    """This function performs a local post-processing of the prediction output,
    operating on each tile generated when the input data is large.
    It is intended for pixel-wise operations, which do not require a global context.

    Parameters
    ----------
        y_pred : numpy array
            tile of the model. It is the result of the prediction, composed of the class probabilities stored in each channel.
            Its shape is
                - (number, width, height, depth=1, nb_channels) in 2D
                - (number=1, width, height, depth, nb_channels) in 3D

    Returns
    -------
        post-processed tile (note: It is currently limited to output a single channel.)
        Its shape is
            - (number, width, height, depth=1, 1) in 2D
            - (number=1, width, height, depth, 1) in 3D
    """
    # This function assumes that the model generates a probability score for each class.
    # It returns the index of the image (label) with the highest probability.
    # It is applied on the tiles of the input data, and thus avoid the allocation
    # of the N scalar arrays representing the probability of each class.
    if len(y_pred.shape) != 5:
        raise ValueError("incorrect 'y_pred' shape, must be a 5d array with shape=(number, width, height, depth, channels)")

    _, _, _, _, channels = y_pred.shape

    if channels == 1:
        local_output = y_pred > 0.5
    else:
        local_output = np.expand_dims(np.argmax(y_pred,axis=-1),axis=-1)

    return local_output


def postprocess_global_output(y_pred, x):
    """This function performs a global post-processing of the prediction output,
    after the tiles have been re-assembled. It is intended to be used for global
    normalization operations.

    Parameters
    ----------
        y_pred : numpy array list
            model prediction output
            Each element of the list has the shape (width, height, depth)
        x : numpy array
            origin data input
            Has a shape (width, height, depth)

    Returns
    -------
        A list of the processed arrays, each with shape (width, height, depth)
    """
    # This function casts the y_pred elements (float32 array) to an 8-bits unsigned integer
    # when handling labels. The resulting image is therefore of this type.
    return [y_pred[i].astype(np.uint8) for i in np.arange(len(y_pred))]


def postprocess_output_type():
    """This function allows specifying the output object type of the Deep Learning Prediction module.

    Returns
    -------
        Object type (i.e. HxUniformLabelField3, HxUniformScalarField3, etc.)
    """
    # This implementation makes the Prediction return a label field dataset,
    # rather than a grayscale image.
    return ['HxUniformLabelField3']

def get_normalization_configuration():
    """Defines the normalization to apply on the input before the prediction.

    Returns
    --------
    A list of normalization configurations: each configuration is defined using a
    CanalConfig(item_or_collection, algoType, per_channel, parameters=None) function.

        item_or_collection: str
            - "PerSample": the normalization is computed for each item of the collection independently
            - "Global": the normalization is computed globally for the whole collection

        algo_type: str
            - "Standardization": applies the normalization (x-mean)/std_dev to the input
            - "MinMax": applies the normalization (x-min)/(max-min) to the input
            - "Percentile": applies the percentile normalization from the [1, 99.8] input percentage range
            - "None": no normalization is applied

        per_channel: bool
            The normalization is performed for each channel or over all channels.

        parameters: DO NOT MODIFY

    If this function is not in the custom file, or the custom file itself is missing, no normalization is performed.
    """
    return [CanalConfig('PerSample', 'Standardization', 'True', parameters=None)]

def get_product_name():
    """Product name.
    DO NOT MODIFY.

    To get the product name using the Python console, type the command
    >>>hx_application.name

    Returns
    -------
        The product name ("Avizo", "Amira" or "PerGeos").
    """
    return "Amira"

def get_product_version():
    """Product version.
    DO NOT MODIFY.

    To get the product version using the Python console, type the command
    >>>hx_application.version

    Returns
    -------
        The product version line number including patch number in format yyyynp.
    """
    return 202220
```
